# Cooperation mitigates diversity loss in a spatially expanding microbial population

**DOI:** 10.1101/668590

**Authors:** Saurabh Gandhi, Kirill S. Korolev, Jeff Gore

## Abstract

The evolution and potentially even the survival of a spatially expanding population depends on its genetic diversity, which can decrease rapidly due to a serial founder effect. The strength of the founder effect is predicted to depend strongly on the details of the growth dynamics. Here, we probe this dependence experimentally using a single microbial species, *Saccharomyces cerevisiae*, expanding in multiple environments that induce varying levels of cooperativity during growth. We observe a drastic reduction in diversity during expansions when yeast grows non-cooperatively on simple sugars, but almost no loss of diversity when cooperation is required to digest complex metabolites. These results are consistent with theoretical expectations. When cells grow independently from each other, the expansion proceeds as a pulled wave driven by the growth at the low-density tip of the expansion front. Such populations lose diversity rapidly because of the strong genetic drift at the expansion edge. In contrast, diversity loss is substantially reduced in pushed waves that arise due to cooperative growth. In such expansions, the low-density tip of the front grows much more slowly and is often reseeded from the genetically diverse population core. Additionally, in both pulled and pushed expansions, we observe a few instances of abrupt changes in allele fractions due to rare fluctuations of the expansion front and show how to distinguish such rapid genetic drift from selective sweeps.

**Significance statement:** Spatially expanding populations lose genetic diversity rapidly because of the repeated bottlenecks formed at the front as a result of the serial founder effect. However, the rate of diversity loss depends on the specifics of the expanding population, such as its growth and dispersal dynamics. We have previously demonstrated that changing the amount of within-species cooperation leads to a qualitative transition in the nature of expansion from pulled (driven by migration at the low density tip) to pushed (driven by migration from the high density region at the front, but behind the tip). Here we demonstrate experimentally that pushed waves, which emerge in the presence of sufficiently strong cooperation, result in strongly reduced genetic drift during range expansions, thus preserving genetic diversity in the newly colonized region.

## Introduction

Spatial population expansions occur at multiple scales, from the growth of bacterial biofilms and tumors to the spread of epidemics across the globe (1–4). Natural populations often undergo range shifts or range expansions, in response to changing climate, and increasingly, following introduction into novel geographical areas due to trade, travel and other anthropogenic factors (5–7). The fate of these spatially expanding populations depends on their genetic diversity, which allows them to adapt to the new environment (8). The very process of spatial expansion is, however, predicted to erode the diversity of the population (9, 10), since the newly colonized territory is seeded by only a subset of the genotypes that exist in the original population. This phenomenon, known as the founder effect, greatly amplifies genetic drift in the population and leads to diversity loss and accumulation of deleterious mutations (11–13). Thus, a firm understanding of the founder effect is necessary to predict and control the fate of expanding species. While diversity is lost during all expansions, the rate of loss is expected to be strongly influenced by the expansion dynamics, which depend on the details of dispersal and growth. Depending on the expansion dynamics, population expansions can be classified into two categories – pulled and pushed. In populations that do not exhibit any within-species cooperation, the growth rate is maximum at low densities and decreases monotonically as the density increases. In such populations, migrants at the low density tip of the wave grow at the fastest rate, and drive the expansion into the new area. Such expansions are called pulled waves, and their expansion velocity, also known as the Fisher velocity, depends solely on the diffusion rate of the individuals and the growth rate of the species at low density. On the other hand, pushed waves occur in the presence of cooperative growth within the population (i.e. positive density dependence of the growth rate, also known as the Allee effect) whereby the tip grows at a much lower rate than the higher density bulk (14–17). Since the growth rate at low density in such populations is lower than in the bulk, the Fisher velocity for such populations is lower than the actual expansion velocity. Although pulled and pushed waves are typically distinguished based on the growth dynamics, a similar distinction can be drawn based on how the dispersal rate depends on the population density (18).

The difference in the dynamics of pulled and pushed waves has substantial genetic consequences (19–21). In its simplest form, range expansions can be viewed as a series of founding events, where a small subpopulation establishes a colony in a new territory, grows rapidly, and then seeds the next founder population. This series of population bottlenecks quickly erodes the genetic diversity in the population, a process aptly called the serial founder effect. The bottlenecks are less severe for species with an Allee effect because growth in the low-density founding colonies is subdued. Indeed, the slow growth of the founders provides sufficient time for the arrival of migrants from the genetically diverse population bulk. Thus, genetic diversity is predicted to persist much longer and over longer distances in populations with an Allee effect (Fig. 1A).

**Figure 1.**
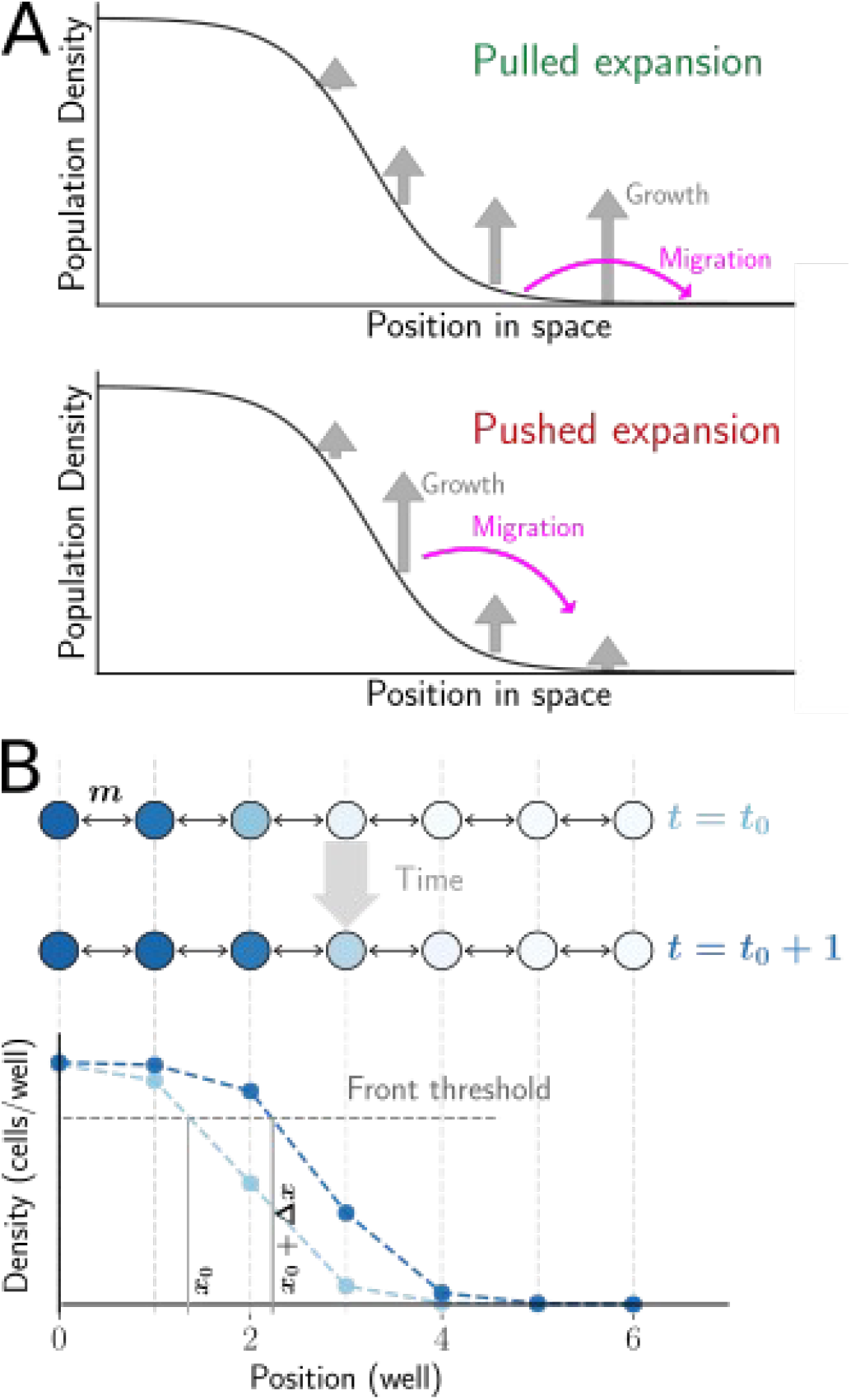
Experimental setup to study genetic consequences of pulled and pushed range expansions. **A.** Range expansions can be broadly classified as pulled or pushed depending on the primary drivers of the expansion. In pulled expansions, the small number of founders from the tip of the expansion grow rapidly in the new territory (top panel). This founding population contains only a small subset of the total diversity in the population. Therefore, diversity is quickly eroded as the population expands into new area. Pushed waves are driven by migration out of the bulk, because the small density of founders at the front has a subdued growth rate (bottom panel). As a result, genetic diversity is maintained much longer. **B.** The experimental setup consists of yeast expanding in a discrete space, discrete time one-dimensional metapopulation landscape. Adjacent wells are connected via migration, and exchange a fixed fraction of cells, m, every cycle, and then grow for 4 hrs (top panel). This process results in an emergent wavefront of a fixed density profile moving to the right with a fixed velocity (bottom panel). The location of the wavefront is determined as the interpolated well position where the density profile crosses a predetermined threshold. Velocity is then measured as the rate of advance of the wavefront location. The entire area to the right of the threshold location is defined as the ‘front’ for subsequent computation of genotype frequencies.

This differential rate of diversity loss in pulled and pushed waves is well-characterized in a wide range of theoretical models (20–24), and has also been observed empirically in field studies (25). However, it has been difficult to directly connect the empirical observations to theory (25), in part because these natural expansions cannot be replicated, and also because numerous environmental factors cannot be well-controlled. Microcosm experiments have helped address this chasm between theory and experiments by partially trading off realism for much better controlled and replicable biological systems (26–30).

Previous experiments with microbial colonies expanding on agar have demonstrated both diversity eroding and diversity preserving range expansions (21, 31, 32). In these experiments, a colony is inoculated with two genotypes, and the diversity loss manifests in the formation and coalescence of monoclonal sectors. However, this sectoring phenomenon is lost when two different mutualist species are inoculated together at the center instead of a single species. The sector formation in the former case and its lack in the mutualists can be well-understood mechanistically for this particular system in terms of the (microscopic) demographic and geometrical properties of the expanding species. In contrast, in our current study, we explore the differential rate of diversity loss more generally as a consequence of growth demographics, independent of species-specific mechanisms. Using the framework of pulled and pushed waves, we performed experiments to establish a general relationship between cooperativity in growth dynamics and the strength of genetic drift. Our setup is an extension of a previously developed experiment, where we demonstrated the transition from pulled to pushed waves with increasing cooperation in yeast (33). To study genetic diversity, we introduced two otherwise identical genotypes with different fluorescent markers, whose frequency can be tracked over time. We find that yeast populations expanding as a pulled wave undergo a drastic reduction in genetic diversity, unlike the same population expanding as a pushed wave. Moreover, we quantify the rate of diversity loss in terms of the effective population size, and show that the effective population size correlates well with how pushed the expansion is (aka ‘pushedness’).

We also observe a few evolutionary jackpot events during which one of the genotypes abruptly increases in frequency. Such events are predicted to arise naturally due to rare stochastic excursions of the expansion front ahead of its expected position (34). Our results support this theory because abrupt changes in allele frequency co-occur with substantial changes in front shape. Importantly, we show that these evolutionary jackpot events can be distinguished from selective sweeps, in which a new mutant rises to high frequency due to its higher fitness than the ancestral population.

## Results

The stepping-stone metapopulation model is widely used to describe the spatiotemporal population dynamics in patchy landscapes (35, 36). In this model, populations grow in discrete patches that are connected to nearest neighbor patches via migration, which is reflected in our experimental setup. The budding yeast, *S. cerevisiae*, expands in one dimension, along the rows of a 96-well plate, with cycles of growth, nearest-neighbor migration, and dilution into fresh media (Fig. 1B). At the beginning of every cycle, a fixed fraction ($*m/2$*) of culture in each well is transferred into wells at adjacent locations on either side, while the remaining ($*1-m$*) is transferred into the well at the same location (migration rate, $*m = 0.4$*,unless stated otherwise). At the same time, the culture is also diluted into fresh media by a constant factor. After dilution, the cultures are allowed to grow for 4 hours before the cycle is repeated. Starting with a steep initial spatial density profile of yeast, this process leads to a stable wavefront (as defined in Fig. 1A, Materials and Methods) that expands at a constant velocity (Fig. 2A).

**Figure 2.**
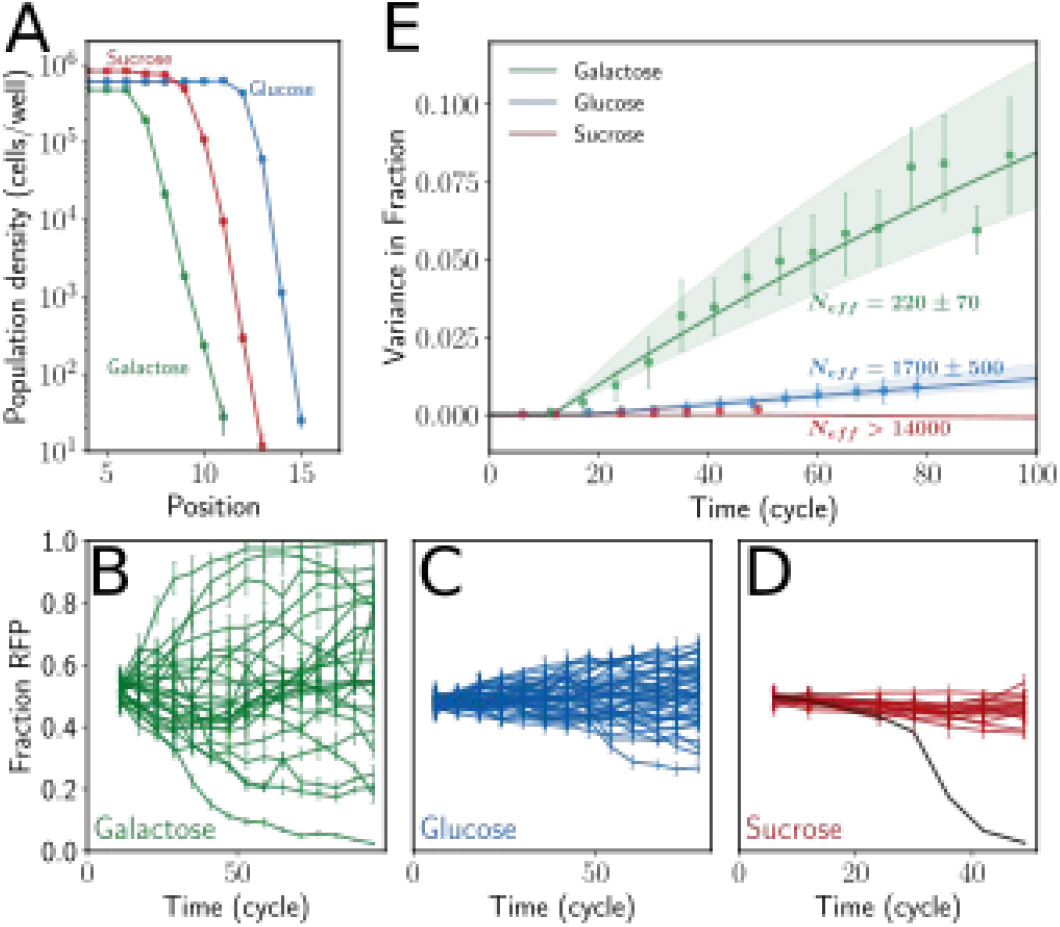
Yeast expanding in different growth media loses diversity at very different rates even though the wavefronts have similar velocity and bulk density. **A.** Populations of *S. cerevisiae* growing in galactose, glucose, or sucrose media expand spatially as traveling waves with a constant velocity and exponentially decaying density at the front. The velocities, bulk population densities, and the shape of the front are similar in all three environments. **B.** Yeast expanding on galactose lose diversity most rapidly. Starting with equal initial frequencies of two genotypes that differ only in terms of a single fluorescent marker (RFP or CFP), the fraction of one of the genotypes in the front (RFP) fluctuates randomly until the genotype either reaches fixation or becomes extinct. The expansion experiments are replicated 24 times, and the dynamics of fractions varies by a large amount across replicates. **C, D.** The same experiments but in different media, glucose and sucrose, show very different rates of diversity loss. In glucose (**C**), the loss of diversity is much slower compared to the expansions in galactose. In sucrose (**D**), no significant loss of diversity is observed during the duration of the experiment(the replicate shown in grey was mis-pipeted in cycle 30, and hence diverges from the rest (SI Fig. 2 top row). This replicate is ignored in further analysis). **E.** The rate of diversity loss can be quantified in terms of the variance between the fractions across (Eqn. 1, f = 0.5). In galactose and glucose, the variance increases significantly, allowing us to quantify the effective population size. In sucrose, the increase in variance is not statistically significant. Thus we can only set a lower bound on the effective population size. The drastic loss of diversity in galactose is reflected in the effective population size of the expanding front, ~220, over four orders of magnitude lower than the actual population size in the front. Effective size in glucose is around 1500, and that in sucrose is estimated to be over 15,000.

Previous studies have shown that yeast typically do not display cooperative behavior when growing on simple sugars such as galactose or glucose, but grow cooperatively on sucrose (37). Thus, we expect pulled expansions in glucose and galactose and pushed expansions is sucrose. To compare the rate of genetic drift in different environments, we use two otherwise identical genotypes of the same strain, but with different constitutively expressed fluorescent markers, whose frequency can be tracked using flow cytometry. We start with a 1:1 ratio of the two strains in the initial density profile for the expansion experiment and observe the relative frequencies for about 100 cycles.

In the galactose environment, the relative frequencies of the two genotypes (as defined in Materials and Methods) change rapidly over the course of the spatial expansion, undergoing large fluctuations, occasionally leading to fixation of one of the genotypes. Twenty four replicate realizations of the experiment reveal that while the waves are nearly identical in terms of their velocity and wavefront shape, the internal dynamics of individual fractions is highly different (Fig. 2B). This can be clearly seen from the variance in fractions across replicates (Fig. 2E), which grows from 0 at the beginning of the experiment to the maximal value of 0.5. The measured variance allows us to quantify the rate of diversity loss in terms of the effective population size using the following relationship:

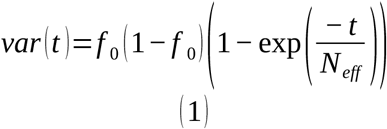

where var(t) is the variance in the fractions across replicates as a function of time, f_0 = 0.5 is the initial fraction at t = 0, and t is in the units of generation time (cycles in this case, since the entire front is effectively diluted by 2x every cycle, and so, each cycle corresponds to one generation). For the pulled waves in galactose, the effective population size is approximately 210 – four orders of magnitude smaller than the actual size of the population in the wavefront (Fig. 2E). We thus see that there is a tremendous loss of genetic diversity during pulled expansions.

We repeat the same experiment, but now with yeast growing on sucrose, where we expect growth to be cooperative and hence, the expansions to be pushed (33). The expansion speed and bulk population density in sucrose is similar to that in galactose (Fig. 2A). Yet, while the waves are physically similar, their effect on the genetic diversity in the population is drastically different. The frequencies of the two genotypes, starting at an equal 1:1 ratio, remain almost unchanged at the end of the experiment (Fig. 2D). The diversity preserving nature of these pushed expansions is reflected in the large effective population size, estimated to be higher than 15,000 – at least two orders of magnitude larger than in pulled waves (Fig. 2E).

Drastically different effective population sizes in simple sugar galactose and complex sugar sucrose are consistent with the theoretical expectations for pulled and pushed waves. Expansions in glucose, however, show somewhat unexpected dynamics. Because glucose is a simple sugar, we expect the expansions to be pulled, and hence lose diversity quickly. However, the measured effective population size in glucose is intermediate between that in galactose and sucrose (Fig. 2C,E), i.e. diversity during glucose expansions is lost much faster than in sucrose, but not quite as rapidly as in galactose.

One possible explanation for this discrepancy is that expansions in glucose are weakly pushed. In order to test this possibility, we quantify the pulled vs. pushed nature of the expansions in all three media. Specifically, we measure the low-density growth rate of our strains and their expansion velocity (DH, see Materials and Methods). Pulled waves expand at the Fisher velocity, which is determined solely by growth rate at low density and the migration rate, while pushed waves expand at a velocity greater than the Fisher velocity. We define a ‘pushedness’ parameter as the ratio of the experimentally observed velocity to the Fisher velocity, so that pushedness = 1 for pulled waves, and > 1 for pushed waves.

For galactose, the pushedness of the expansions is observed to be close to 1, whereas that for sucrose is 2.3, clearly confirming that the galactose expansions are pulled and the sucrose ones are pushed (Fig. 3A). Surprisingly, the pushedness for glucose expansions is also greater than 1, suggesting that contrary to our naïve expectation, expansions in glucose are in fact not pulled. More careful measurements of the growth profile of the DH strains in 0.2\% glucose reveal a very tiny amount of cooperative growth at extremely low densities (below $10^3 cells/well$), making them very weakly pushed (SI Fig. 1). While this Allee effect might originate due to many possible factors such as collective pH modulation (38), it is important to note that the emergent property of the wave, pushedness, explains the decreased rate of diversity loss without the need to understand species-specific growth mechanisms.

**Figure 3.**
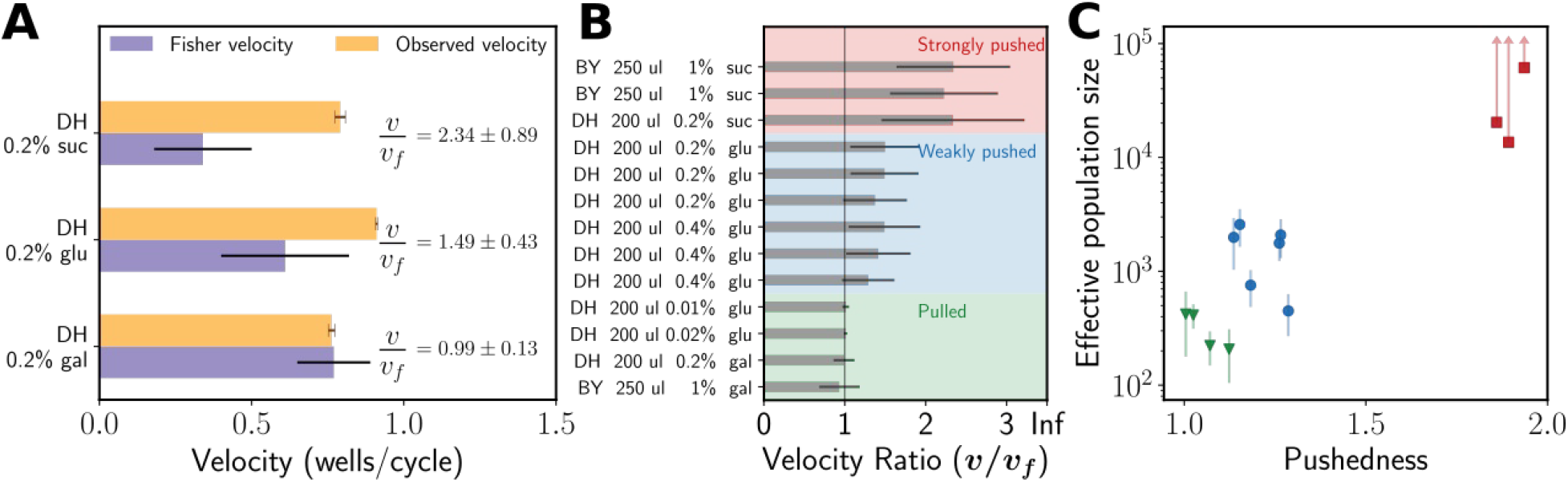
The ratio of observed velocity to the Fisher velocity (termed pushedness) determines the rate of diversity loss during expansions. **A.** Pulled waves expand at the Fisher velocity, and have a pushedness of 1, whereas pushed waves have pushedness larger than 1. Consistent with the observed rates of diversity loss, the waves in galactose have pushedness = 1, and those in sucrose have a much large pushedness of 2.3. Even though digestion of glucose is non-cooperative, we found expansions in glucose to also be pushed although more slightly than in sucrose. This explains the intermediate rate of diversity loss in glucose compared to galactose and sucrose. **B.** We repeat the expansion experiments across multiple environmental conditions (media, death rate, migration rate), for two different pairs of yeast strains (BY and DH) and observe a wide range of pushedness values for the different expansions. **C.** Effective population size is plotted against the pushedness for the different strain-media combinations. We find that N_{eff} correlates strongly with the pushedness (note the log scale).

We further probe the relationship between pushedness and the rate of diversity loss experimentally, by repeating the expansion experiments in multiple environments using two different pairs of strains (DH-RFP/DH-CFP and BY-RFP/BY-YFP). The different strain-media combinations give rise to expansions spanning a broad range of pushedness values (Fig. 3B). We find that the pushedness correlates well with the effective population size during expansions (Fig. 3C). Broadly, for all instances of pulled waves, Neff was under 500, over four orders of magnitude below the actual population size. Within the pushed waves, we find two regimes with very different rates of diversity loss. In the weakly pushed regime, the effective population size ranged between 500 and 4000. We thus see that even for pushed waves, if the cooperativity is not strong enough, diversity can be lost quite rapidly. Finally, in the strongly pushed regime, we observe very little genetic drift and can therefore only set a lower bound on the effective population sizes (Materials and Methods), and the lower bounds are at or over 15,000 (Fig. 3C). Overall, for populations with approximately equal bulk densities (within a factor of 3), the rate of diversity loss is seen to be strongly modulated by the pushedness.

Throughout our experiments, we observe a few instances where one of the genotypes appears to take over the population very rapidly. Fig. 4A shows two such rapid takeover events, which closely resemble evolutionary sweeps. However, during range expansions, such sweeps can also occur purely as a consequence of a rare reproduction or dispersal event. In the wild, a rare long-distance dispersal might establish a new population in an unoccupied territory near the front. When this nearly clonal population merges with the expanding front, the frequency of the dominant genotype in the front suddenly increases. This process, called the ‘embolism effect’ has been previously proposed in theoretical literature (24), and we found one instance of it in our experiments (SI Fig 2 top panel). In our experiments, rapid takeovers might also occur when a clump of cells of the single genotype is transferred over to the front of the wave, leading to increased frequency of that genotype in the front. As the expansion progresses, this increased frequency propagates through the entire front (Fig. 4B, top panel). Both examples above can be termed a jackpot event that occurs due to stochastic demography.

**Figure 4.**
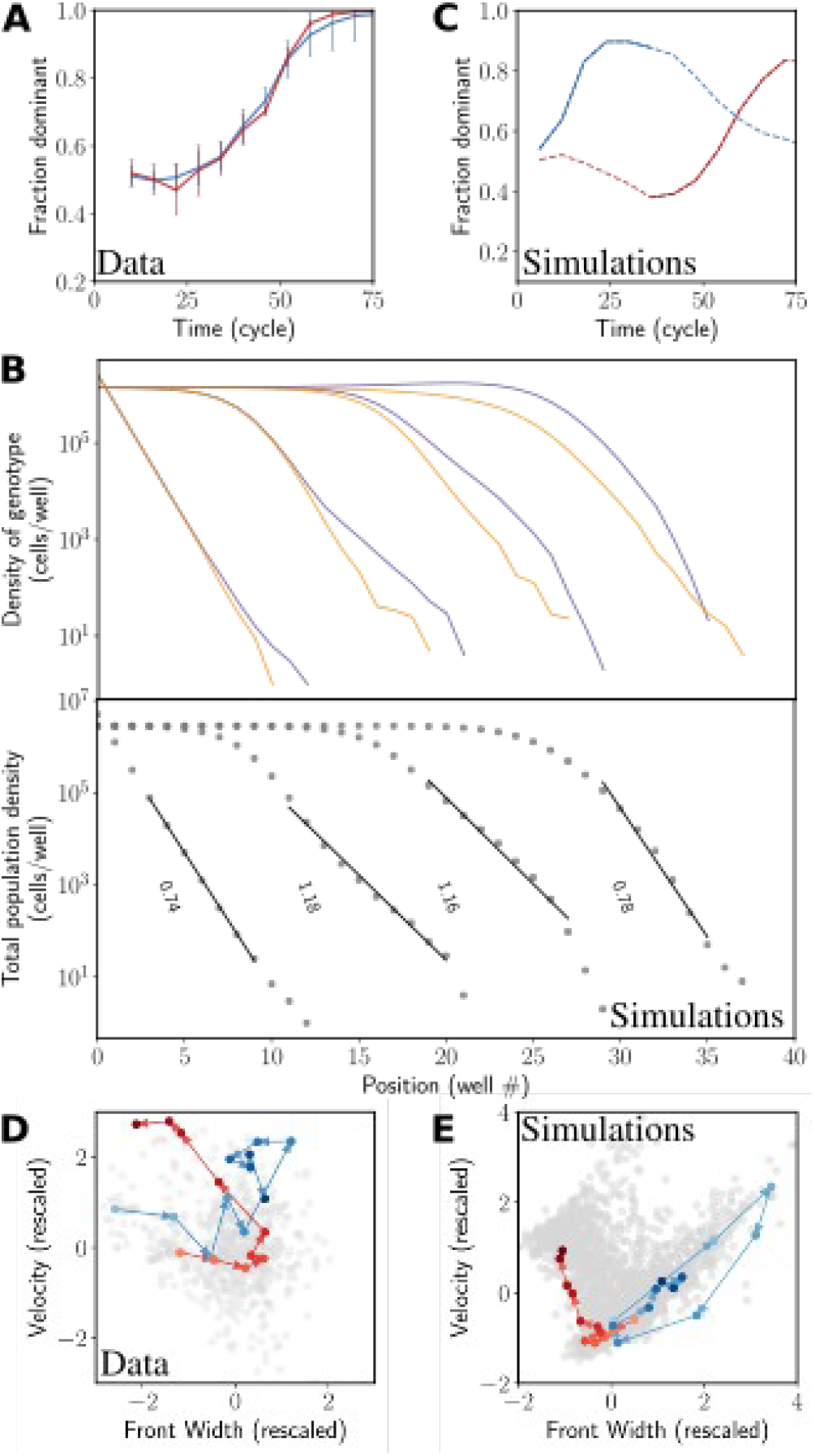
Rapid takeover by one of the genotypes due to rare fluctuations of the front and selective sweeps. **A.**In some instances of the expansion experiments, the fraction of one of the species is seen to increase very rapidly. The fraction of the species that eventually dominates is plotted as a function of time for two such instances. **B.**During spatial expansions, rapid takeovers can occur without any selection, simply as a result of stochasticity in migration and growth, or rare long distance dispersal. The top panel shows the density of two genotypes in a simulation at different times. In an early cycle, at the very tip, stochasticity in migration led to excess colonization of the purple genotype in a well near the front (jackpot event). This fluctuation then propagated back towards the bulk as the purple genotype rapidly took over the front. Note how this process was accompanied by a transient widening of the front (bottom panel). **C.** Two instances of rapid takeovers in simulations. The orange curve is from a simulation of a selective sweep during expansion, whereas the blue curve corresponds to a jackpot event. The dotted lines are the entire trajectory, and the solid sections correspond to the takeover times that are analyzed further. **D, E.** Trajectories in the space of front width and velocity for experiments (**D**) and simulations (**E**) from **A** and **B**. Each dot corresponds to the front width and velocity at a single time point for one of the replicates. The axes are rescaled so that the front width and velocity have mean 0 and standard deviation of 1; arrows indicate increasing times. During selective sweep (orange curve in **E**), the trajectory initially fluctuates around the mean value of the width and velocity, but, after the mutant establishes at the front, the trajectory moves monotonically to the top left towards increasing velocity and decreasing front width. In contrast, for the jackpot event (blue), the front width and velocity transiently increase, but relaxe back towards their mean values at later times. Although the timeseries of the fractions in experiments looks nearly identical in the two instances shown, the state space trajectories are clearly distinct.

A completely different possibility is that a rapid increase in the frequency of a neutral allele is due to a selective sweep due to a mutation at another locus. We can distinguish the two via the excess migration at the front that accompanies jackpots but not selective sweeps. In Fig. 4B, we simulate a simple model of expansion to show how the wave front widens as a consequence of the excess migration. Wider fronts expand faster, so the wave speed increases transiently as well. Importantly, both the velocity and front width return to their mean values as the front returns to equilibrium. In contrast, evolution towards a higher growth rate (migration rate is fixed in our assay and cannot be selected for) leads to increased velocity, but decreased front width (front width of pulled waves $\sim\frac{1}{\ sqrt{r_0}}$, (14)). Moreover, in the case of selective sweeps, the trajectories in the velocity-front width space do not return to the previous mean, but rather settle at the new equilibrium. These differences allow us to distinguish between the two processes responsible for rapid takeover by a genotype.

The dynamics described above are confirmed in simulations, where, we follow the trajectory of a rapid takeover event in the state space (front width – velocity). For jackpots (no beneficial mutations allowed), we see the transient front widening accompanied by an increased velocity, before the trajectory returns to the mean front width and velocity (Fig. 4E). When a low rate for benefitial mutation rate is included in our simulations, we observe rapid extinctions of one of the neutral markers. The state space trajectories are, however, very different. After a selective sweep, they do not return to their previous locations; instead, they settle in the region of higher velocity and steeper fronts (Fig. 4E).

Among the rapid takeovers that we observe in experiments, a subset can be clearly seen to follow the selection template. Fig. 4D shows the state space trajectory for one replicate that putatively evolved to a higher growth rate (red trajectory, compare to a jackpot shown in blue), corresponding to the takeover trajectories shown in Fig. 4A. We observe these putative selective sweeps only in a single growth medium among several that we used in our experiments (SI Fig. 3). This medium was limiting in terms of an essential amino acid, and thus, is likely to apply a higher evolutionary pressure than the others. We also observe a few rapid takeover events that do not follow the selection template, but rather, look like jackpot events. Even though the time series of allele fraction look similar for selective sweeps and jackpot events (Fig. 4A), the two mechanisms can be clearly distinguished based on their state space trajectories (Fig. 4D). Given the rarity of both jackpots and selective sweeps due to mutation, we do not have sufficient data to explore and contrast them in great quantitative detail. The few instances of these processes that we do observe are nevertheless fully consistent with theoretical predictions and our simulations.

## Discussion

In this study, we used a well-controlled laboratory microcosm setup to probe the distinct evolutionary consequences of pulled and pushed expansions. We observed the rapid loss of diversity due to the serial founder effect when yeast expanded as a pulled wave, and a much more subdued loss of diversity when it expanded as a pushed wave. Moreover, we explored environmental conditions that span different levels of pushedness and saw found that the effective population size in the front is strongly correlated with the pushedness of the expansion. Thus, our experiments suggest that pushedness is a useful measure for predicting the rate of diversity loss during range expansions.

We also observed instances of unusually rapid take over by one of the genotypes. In the amino-acid-limited media, the yeast evolved a higher growth rate, and the takeover events were driven by selective sweeps. In other conditions, rapid takeovers were instead due to rare demographic fluctuations. We were able to distinguish the two by looking at the trajectories of the wavefronts in the state space defined by velocity and front width.

The extensive theoretical work on range expansions has led to other very interesting predictions that could also be addressed using our experimental system. One prediction pertains to the quantitative dependence of the effective population size on the actual population size of the wavefront (23). It has been established that, with growth and migration held fixed, $N_{eff}$ scales linearly with $N_{bulk} $ in fully-pushed expansions, and $N_{eff}\sim log^3\left(N_{bulk}\right)$ in pulled expansions. Moreover, in the presence of a very weak Allee effect, Birzu et al predict a third class of expansions that is intermediate between pulled and pushed, where $N_{eff}$ scales with $N_{bulk}$ as a sublinear power. We made an attempt to observe these different scaling relationships by varying the bulk population size in experiments in two different ways – by changing the total volume, and thus the population size, and by changing the amount of a limiting amino acid. Unfortunately, in the former case, the altered volume also altered the density-dependence of the growth, while in the latter case, the low amino acid condition led to evolution during expansion. We speculate that the expansions in glucose, where the loss of diversity is intermediate between galactose and sucrose, might in fact belong to the newly predicted third class of expansions. Modifying our assay to modulate the bulk density without changing growth properties would help resolve this speculation.

Demographic stochasticity and environmental noise have also been predicted to cause fluctuations in the position of the expansion front (23, 34), which are well described by simple diffusion. In pushed waves, the effects of demographic noise on front diffusion are predicted to be subdued, and front diffusion should largely reflect the environmental noise. The situation is different in pulled waves, where front diffusion due to demographic noise is predicted to be much more pronounced. We observed front diffusion in both pulled and pushed waves in our experiments, where the variance in front position remains constant for some initial period before it starts increasing linearly with time (SI Fig. 4). Contrary to expectations, we do not find a significant quantitative difference in front diffusion in pulled vs. pushed waves. This negative result could be explained by the lack of a sufficiently long timeseries data or by the dominance of the environmental noise for both pulled and pushed expansions in our experimental setup.

Allee effects, or the inability of organisms to grow optimally at very low densities, is often considered to have a negative impact on populations. For instance, it leads to lower expansions velocities compared to the velocity if growth were not suppressed at the low density tip. However, in this study we demonstrate that the Allee effect can in fact have a very beneficial effect on the expanding population by helping preserve diversity as the population enters novel territories, where the diversity is especially critical for survival. Even a miniscule Allee effect at very low densities, such as we found in the glucose expansions, can go a long way in helping mitigate diversity loss. Perhaps such tiny Allee effects pervasive in many invading species explain the lower than predicted rates of diversity loss during their expansion.

## Materials and methods

### Strains

The expansion experiments were performed using two pairs of strains, BY-RFP/BY-YFP and DH-RFP/DH-CFP. The BY strains were derived from the haploid BY4741 strain (mating type \textbf{a}, EUROSCARF, (39)). The BY-YFP strain has a yellow fluorescent protein expressed constitutively by the ADH1 promoter (inserted using plasmid pRS401 containing MET17). The BY-RFP strain has a red fluorescent protein inserted into the HIS3 gene using plasmid pRS303. The DH strains are the same as those used in Healey et al (40). They are derived from the diploid strain W303, with the RFP/CFP strains harboring constitutively expressed fluorescent markers integrated into the URA3 gene. This pair is auxotrophic to uracil.

### Growth rate measurements and calculation of Fisher velocities

Growth rates for both strain pairs were measured independently for all media, in growth conditions identical to the final expansion experiments. For each pair, the two fluorescent strains were mixed in 1:1 ratio in log phase and the cultures were diluted into a wide range ($10 cells/well$ to $10^5 cells/well$) of initial cell densities. They were then diluted 2x every 4 hours into fresh media. Initial and final densities of each fluorescent strain for each dilution cycle were measured using flow cytometry, and their growth rates as a function of cell density were derived from these measurements. The data is shown in SI Fig. 1. Low density growth rates were obtained by linear regression on the log of initial and final densities, for initial densities under $500 cells/well$. The Fisher velocities were then derived by simulating expansions with logistic growth, with the fitted low density growth rate. Uncertainty in Fisher velocities was obtained by bootstrapping.

### Expansion experiments

All experiments were performed at 30 \degree C in standard synthetic media (yeast nitrogen base and complete supplement mixture), in $200-\mu L$ batch culture in BD Biosciences Falcon 96-well Microtest plates. Expansions occurred along the 12 well long rows of the plate. Migrations and dilutions were performed every 4 h using the Tecan Freedom EVO 100 robot. Plates were not shaken during growth. Optical densities were measured on the robot before every dilution cycle in the Tecan Sunrise platereader with 600-nm light. Cell densities of individual fluorescent strains were also measured every 6 cycles in the MacsQuant flow cytometer after dilution in phosphate buffered saline (PBS). All expansions started with a steep exponential initial density profile. Periodically during the expansion, the leftmost well (in the bulk of the wave, away from the wavefront) was discarded and the entire profile was shifted to the left, so as to create empty wells for further expansion to the right. It was ensured that the rightmost two wells were always at zero cell density so as to avoid any edge effects on the expansion.

### Definition of front

The ‘front’ is defined as the region of the wave density profile that falls below a threshold density, set at $0.2\times N_{bulk}$. ‘Fractions/frequency in the front’ correspond to the fraction of red or green fluorescent cells added up over the entire front region as defined above. The location of the front is defined as the interpolated well position where the density profile crosses the threshold.

### Lower bound on effective population size

Equation 1, which quantifies the dependence of the effective population size on the rate at which variance in fractions across replicates increases, is used to estimate the effective population size in our analysis. However, for pushed expansions in sucrose, the variance in the measured fractions never increases significantly above 0 given the uncertainties in fraction estimation. In this case, it is not possible to actually estimate the effective population size. However, the fact that after a given time T, the variance increases at most by the amount equal to the measurement uncertainty, V_{min}, sets a lower bound on the effective population size:

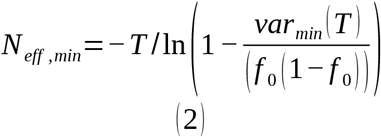

## Supplementary Information

**S1 Fig. 1:**
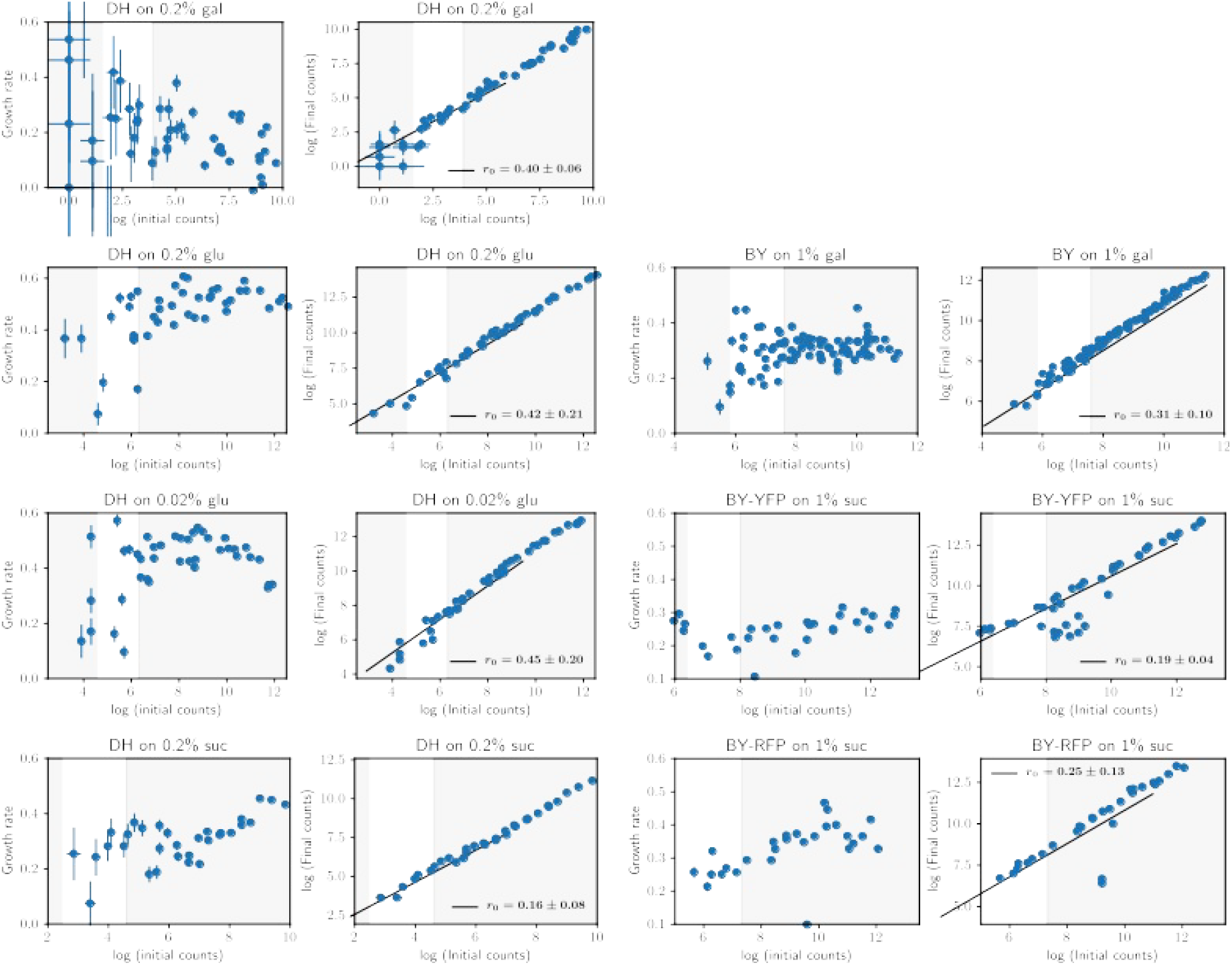
Density dependence of growth rate and fits for low density growth rate for different strain-media combinations. Seven pairs of plots are shown for 7 different strain-media combinations. In each pair, the left plot shows the instantaneous growth rate from each measurement of initial and final densities over a 4 hr period. The right panel shows the raw initial and final densities at the beginning and end of the 4 hr periods on a log scale. Growth rate can be estimated from the right panels by fitting a straight line over the region of interest (shown in white). The fitting region is chosen so that only growth at low density is considered (actual density < ~2000 cells/well). We also exclude very low density data (measured density < ~50 cells/well) from fitting because of the very high sampling noise introduced when measuring at extremely low densities (note that the actual cell density is obtained by multiplying the measured cell density by the dilution factor used for measurement; cells need to be diluted into a buffer because very high densities cannot be measured in the flow cytometer, and it is difficult to have different dilution factors for each well). Uncertainties in growth rates are obtained by bootstrapping.

**SI Fig. 2:**
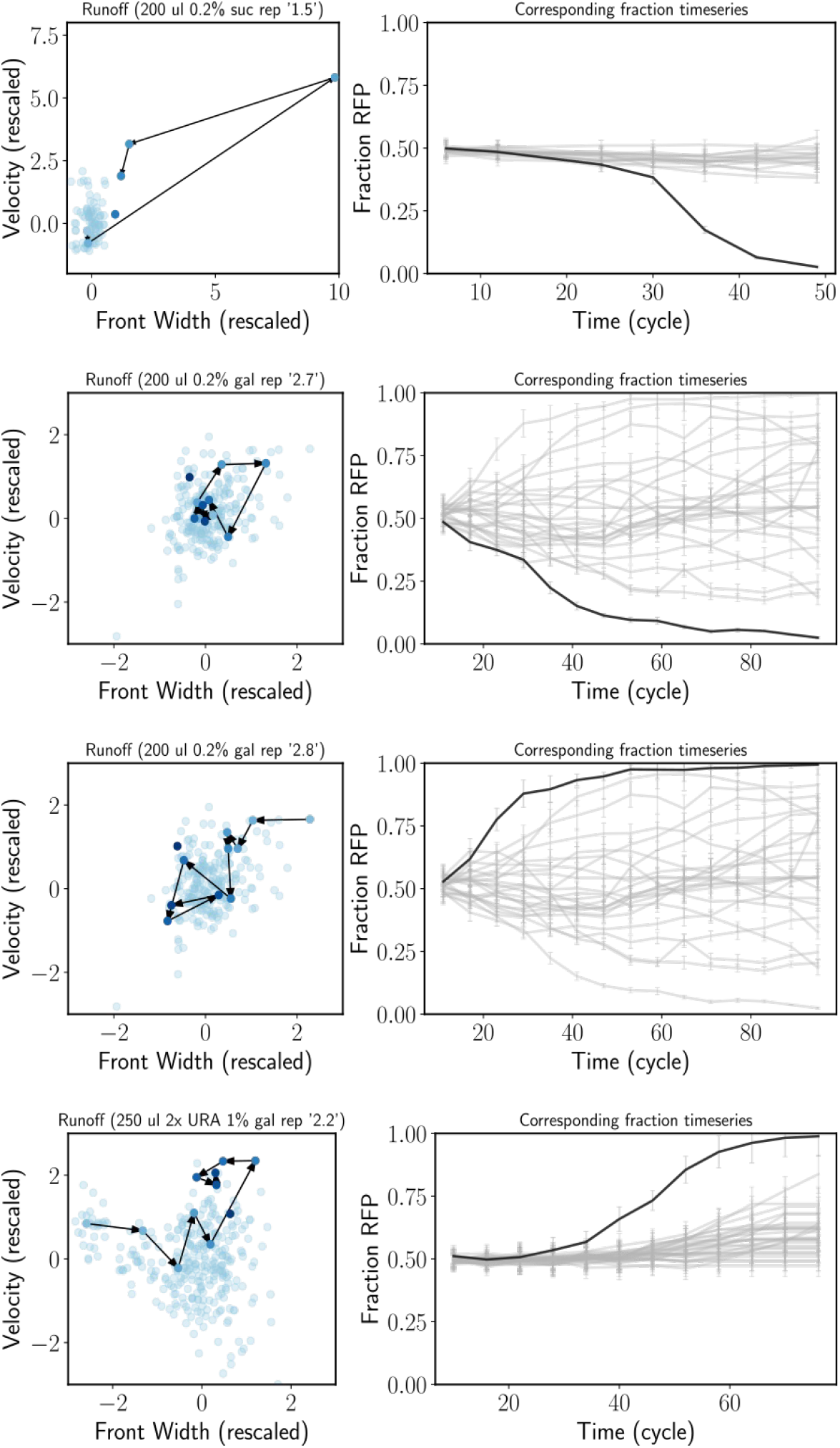
Jackpots in experiments. Jackpots are rare events where stochasticity in migration leads to the front elongating at the very tip, leading to rapid change of fractions of the genotypes. The figure shows 4 instances of jackpots in our experiments. Left panel shows the trajectory of the experiment in the front width-velocity state space. For jackpots, such trajectories transiently move to (or start from) the top right but eventually move back towards the mean (transient elongation of the front accompanied by increased velocity). The right panels show the specific replicate that underwent a jackpot event, compared to the rest of the replicates in the same experiment.

**SI Fig. 3:**
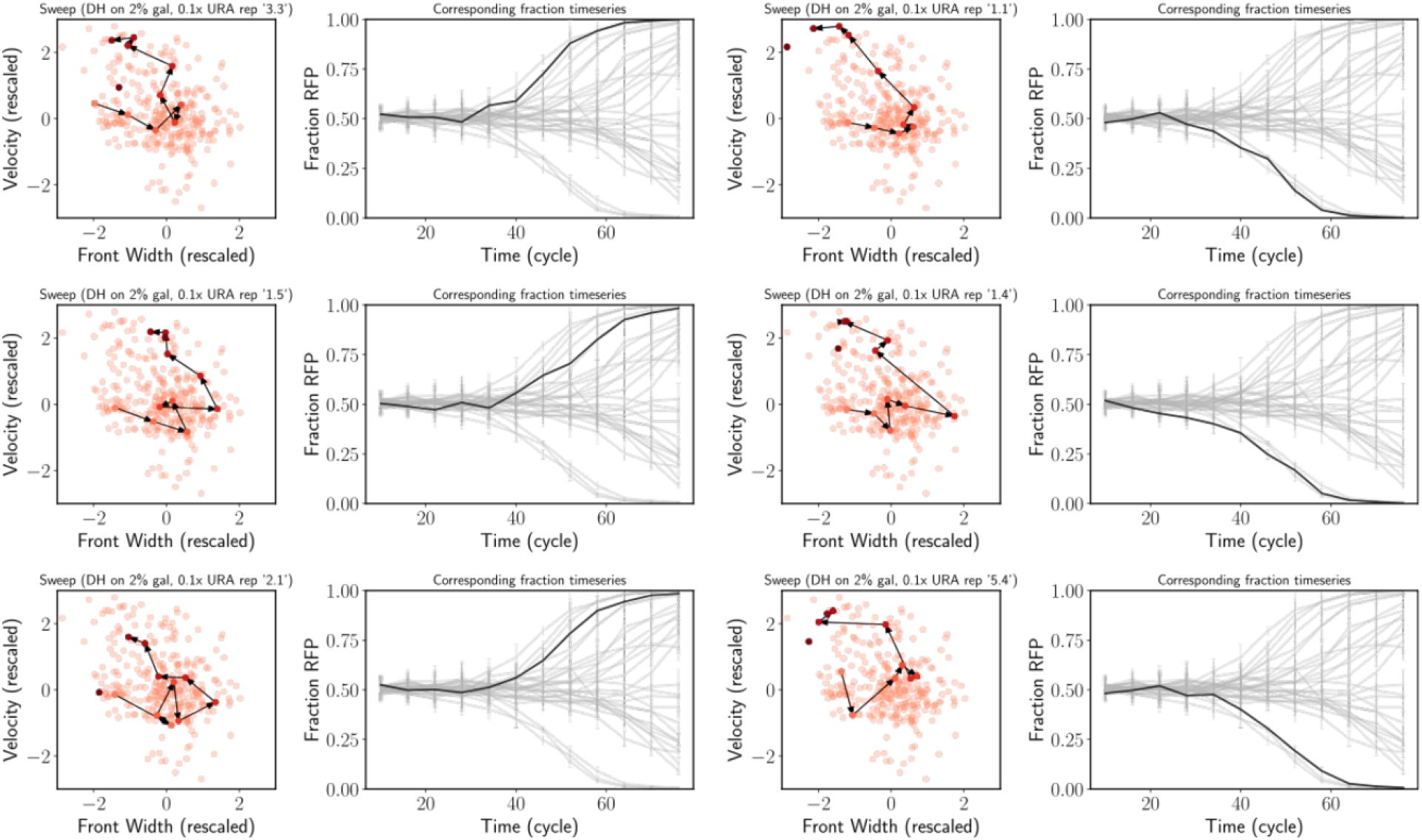
All evolution state space diagrams. Similar to jackpot events, when a faster growing mutant appears in the front, the mutant fraction again increases quickly. However, in contrast to jackpots, where the state space trajectories relax back to equilibrium values, for selective sweeps, the front width decreases and the velocity increases permanently, leading to trajectories that move to the top left in the state space diagram and stay there. The figure shows 6 instances of selective sweeps observed in our experiments.

**SI Fig. 4:**
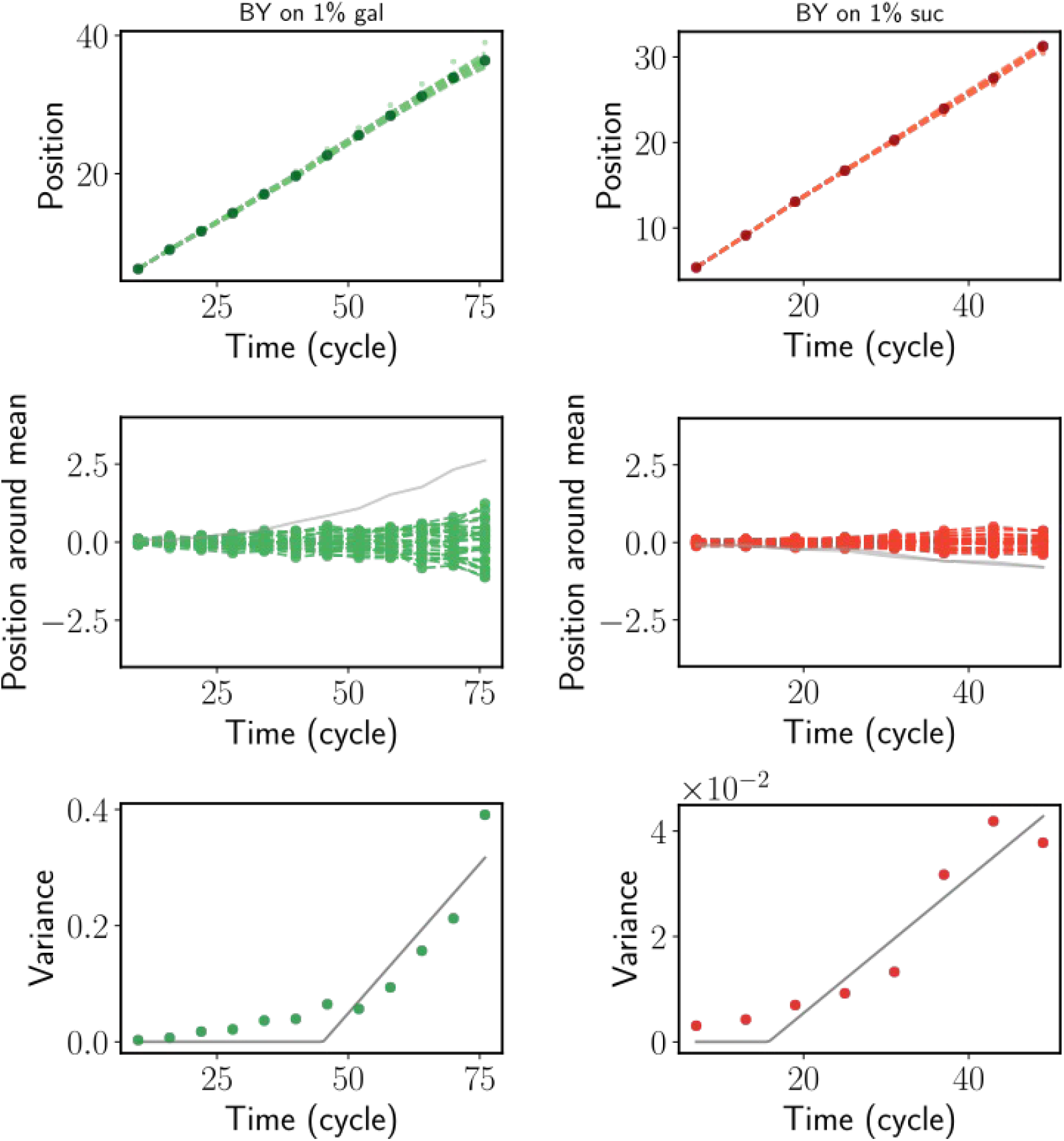
Front diffusion figure. The expansion wavefront is known to diffuse around its mean position as the population expands. The diffusion coefficient is determined by the intrinsic demographic stochasticity in the population as well as the environmental noise. Pulled waves are predicted to be dominated by intrinsic stochasticity, and typically diffuse more around the mean compared to pushed waves, where front diffusion is predicted to be dominated by environmental noise. However, in our experiments, we did not find a significant difference in pulled and pushed waves, suggesting that the diffusion of the front is dominated by environmental noise in both cases, induced by the sampling noise in migration and dilution.

**SI Table 1:**
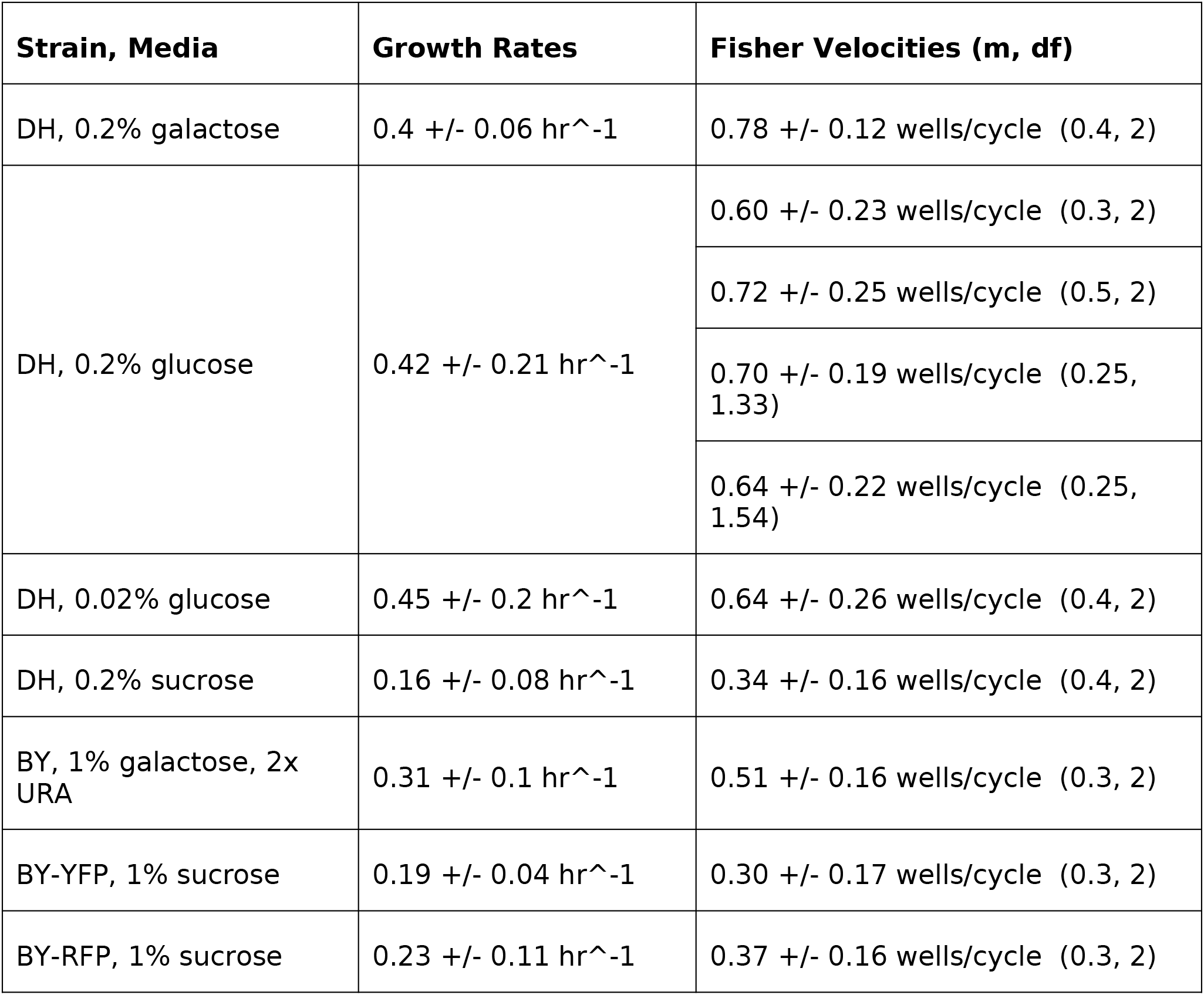
Growth rates and Fisher velocities for different strain-media combinations

